# Pan- and core- network analysis of co-expression genes in a model plant

**DOI:** 10.1101/051656

**Authors:** Fei He, Sergei Maslov

## Abstract

Genome-wide gene expression experiments have been performed using the model plant Arabidopsis during the last decade. Some studies involved construction of coexpression networks, a popular technique used to identify groups of co-regulated genes, to infer unknown gene functions. One approach is to construct a single coexpression network by combining multiple expression datasets generated in different labs. We advocate a complementary approach in which we construct a large collection of 134 coexpression networks based on expression datasets reported in individual publications. To this end we reanalyzed public expression data. To describe this collection of networks we introduced concepts of ‘pan-network’ and ‘core-network’ representing union and intersection between a sizeable fractions of individual networks, respectively. We showed that these two types of networks are different both in terms of their topology and biological function of interacting genes. For example, the modules of the pan-network are enriched in regulatory and signaling functions, while the modules of the core-network tend to include components of large macromolecular complexes such as ribosomes and photosynthetic machinery. Our analysis is aimed to help the plant research community to better explore the information contained within the existing vast collection of gene expression data in Arabidopsis.

**Research Area:** Systems and Synthetic Biology

**Summary:** By analyzing 134 microarray datasets for Arabidopsis, we found that gene coexpression networks are highly context-dependent.

**Financial source:** Work at Brookhaven was supported by grants PM-031 from the Office of Biological Research of the U.S. Department of Energy.

## INTRODUCTION

Coexpression networks represent all pairwise relationships of genes that have similar profiles in a given set of expression samples. Expression levels of such connected genes are either directly or indirectly co-regulated by the same regulatory elements. Since genes under the same regulatory control tend to be functionally associated, the most common use of coexpression networks is to infer unknown and to validate known gene functional roles and regulatory interactions between genes (Kim et al., 2001). A coexpression network consists of all significantly correlated pairs of genes with correlation coefficients above a certain threshold. Once a coexpression network is constructed, the next step usually involves identification of densely interconnected modules, which are often enriched with genes involved in a specific biological function (Stuart et al., 2003). Coexpression network analysis has been actively used in plant functional genomics. For example, new genes involved in flavonoid biosynthetic process (Yonekura-Sakakibara et al., 2007), starch metabolism (Mentzen et al., 2008), aliphatic glucosinolate biosynthesis (Gigolashvili et al., 2009), lignin biosynthesis (Alejandro et al., 2012; Vanholme et al., 2013) and photorespiration (Pick et al., 2013) have been identified with the help of coexpression networks. Compared with animal (especially human) data, functional gene annotation in plants is less comprehensive even in a well-studied model organism such as Arabidopsis. Thus it is especially valuable to leverage the use the existing plant transciptomics data in order to improve identification of new and validation of existing gene and protein functions (Berardini et al., 2004; Lee et al., 2010; Hwang et al., 2011; Heyndrickx and Vandepoele, 2012). Pan-and core-coexpression network analysis proposed in this study serves exactly this purpose.

A common approach to building coexpression networks is to infer correlation relationships from a combination of multiple expression datasets produced by different labs (Kim et al., 2001; Stuart et al., 2003; Atias et al., 2009; Mao et al., 2009; Wang et al., 2012). For example, *Mao et al*. combined all the datasets from the AtGenExpress project (~1000 samples) to construct a coexpression network (Mao et al., 2009). The larger sample size improves the statistical significance of relationships between genes. The inevitable experimental noise within microarray data may give rise to false positive interactions in which pairs of genes have high degree of coexpression in only one dataset but very low coexpression in other datasets (Lee et al., 2004). The traditional approach relies on increasing the number of samples to infer more reliable correlation relationships (Weirauch, 2011). On the other hand, indiscriminately combining multiple samples may not be universally good. The combined samples need to be biologically comparable (Ramasamy et al., 2008). Furthermore, batch effects may give rise to false positive and spurious correlations between genes when microarray data from different labs are combined (Fare et al., 2003; Chen et al., 2011).

Another problem with combined co-expression networks is that it may miss rare gene interactions formed under specific conditions such as a particular disease (de la Fuente, 2010). Increasing amount of evidence indicates that different gene networks operate in different biological contexts (Roguev et al., 2008; Bandyopadhyay et al., 2010). Thus, it is increasingly important to compare and contrast coexpression networks generated from individual datasets (Choi et al., 2005; Ideker and Krogan, 2012; Amar et al., 2013). Experimental results suggest that more than one third of genetic interactions are condition-specific (Guénolé et al., 2013). Several studies have also demonstrated that coexpression of genes varies in different conditions. *Southworth et al*. studied the difference between coexpression networks of young and old mouse brains and found genes involved in memory have more network connections in the young than in the old animals (Southworth et al., 2009). By leveraging the concept of differential rewiring, *Hudson et al*. captured the phenotypic differences between two breeds of cows (Hudson et al., 2009). Compared with normal tissue, many coexpression relationships were lost in cancer tissue, implying the loss of correlated regulation of pathways (Anglani et al., 2014).

A published study of changes in gene expression usually has its own experimental design created in order to answer a specific biological question or several related questions, such as, to understand the mechanisms of plant heat shock response (Charng et al., 2007) or biological function of a plant hormone (Okushima et al., 2005). Here we construct and analyze a comprehensive collection of 134 coexpression networks each based on expression samples from an individual published study, thus preserving context-specific network structure. We assume the network generated from each Gene Expression Omnibus (GEO) series represent a particular regulatory response specific to the experimental design and biological query of that study. Therefore, expression datasets from individual studies are ideal for the detection of condition-specific networks. In this study, we calculated and analyzed the coexpression networks for each of the 134 public microarray datasets in NCBI Gene Expression Omnibus (GEO) database which passed our filters on the minimal number of samples and specific microarray technology.

## RESULTS

### A large collection of Arabidopsis coexpression networks

Previous work has combined expression datasets from many labs to build coexpression networks for animals or plants (Kim et al., 2001; Atias et al., 2009; Mao et al., 2009). Although this strategy has been intensively used by plant scientists to assist gene characterization (Usadel et al., 2009), it preferentially capture the relationships between genes that are conserved across most contexts. In contrast to this, in this study we built coexpression networks based on individual expression datasets from the model plant Arabidopsis in order to capture network aspects that appeared in a specific experimental setup (de la Fuente, 2010).

Thousands of gene expression profiling datasets are available for Arabidopsis in public repositories such as GEO. We focused on the Affymetrix GPL198 platform, since it is the most widely used platform and its annotation is continuously updated. Many of those datasets are not suitable for network inference simply because the number of samples are not enough for a robust inference of correlation. Similar to a previous study performed in humans, we limited our analysis to datasets with at least 20 samples; 134 such datasets were acquired from GEO. Each of them was normalized in the same manner, and genes with very low mRNA abundance or genes without any significant changes were removed (see Methods). The top 0.1% most coexpressed gene pairs within each dataset were then used to build individual networks (Bergmann et al., 2004). We used GSE series number to identify individual published studies, thus each of our 134 networks is labelled with the GSE number of the corresponding GEO experiment/series (Figure 1).

**Figure 1.**
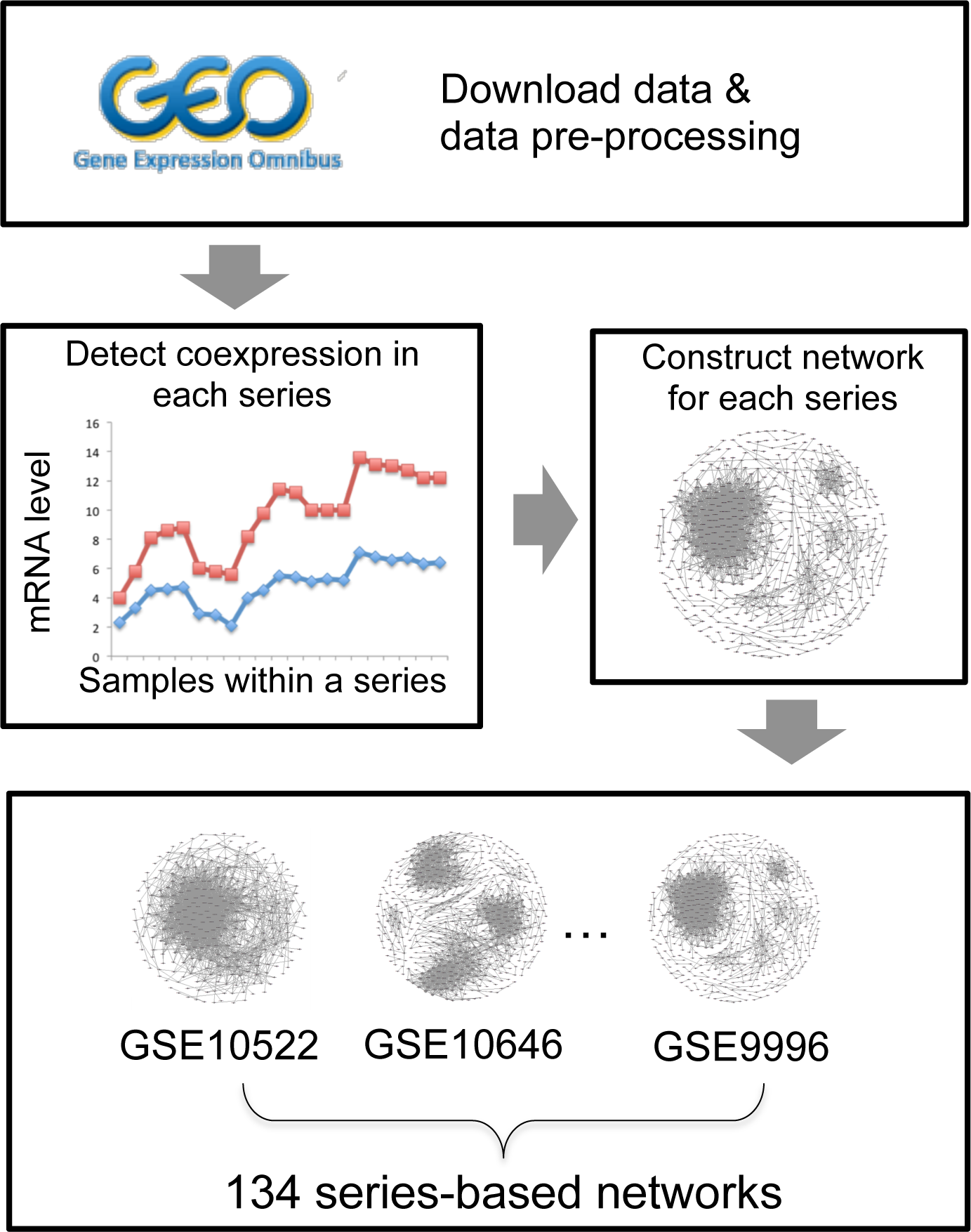
Our pipeline for construction of 134 GSE series-based coexpression networks in Arabidopsis.

The number of nodes in most of the series-based networks was between 500 and 5,000, while the number of edges was between 50,000 and 100,000 (Figure 2). The lowest Pearson Correlation Coefficient (PCC) cutoff for each series-based network was about 0.73 (Figure 2), which is statistically significant for 20 samples (*p-value* < 0.001). About 60% of these series-based networks were inferred from leaf, seedling and root tissues (Figure 2). The other 40% included other tissues such as flowers, seeds and shoots (Figure 2). As can be seen in Figure 2, these series-based networks were inferred from samples in a variety of experimental contexts, such as the effect of gene mutations or abiotic stress. The breadth of data sources were included to better capture of coexpression networks in different conditions.

**Figure 2.**
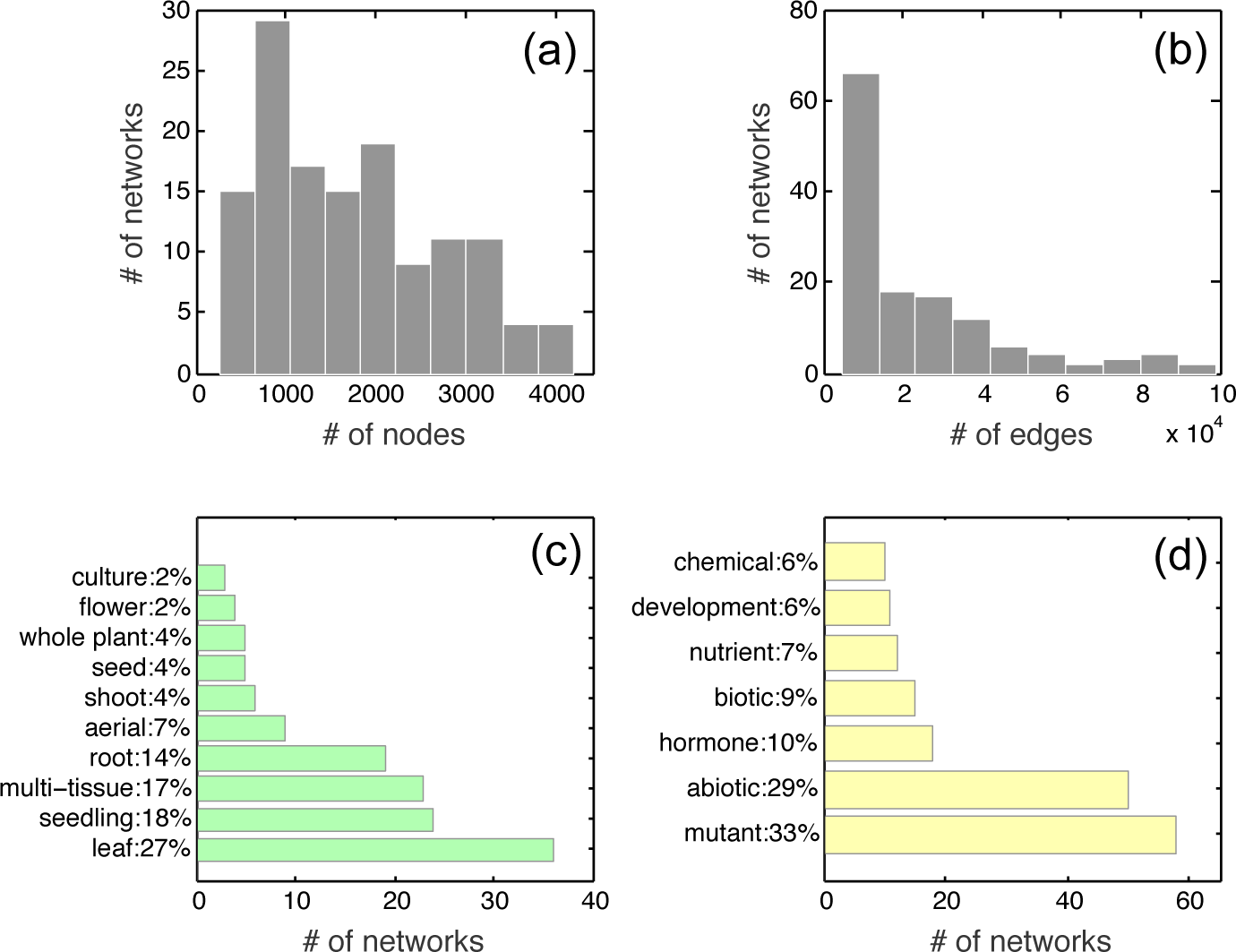
Basic statistics for 134 GSE series-based networks. (a) The histogram for the number of nodes in each series-based network; (b) The histogram for the number edges in each series-based network; (c) The bar chart of tissue types of the data source; (d) The bar chart of experimental conditions of the data source.

### Coexpression networks in Arabidopsis are highly context-dependent

Similar to the idea of ‘pan/core genome’ in bacteria (Medini et al., 2005; Lapierre and Gogarten, 2009) and plants (Hansey et al., 2012; Hirsch et al., 2014; Golicz et al., 2015), we propose to use the term ‘pan-network’ for the union of all the 134 series-based networks, while ‘core-network’ to represents the intersection of a considerable fraction (10 or more out of 134) of these networks. Indeed, unlike a large number of core genes shared across the entire there were no edges present in all 134 of our networks. This is similar in spirit to a soft cutoff used by one of us (Dixit et al., 2015) in detecting the core (“basic”) genome of a bacterial species (*E. coli*).

The pan-network representing the union of all 134 of our individual networks contains 2,294,175 non-redundant edges and 18301 nodes/genes (Supplemental File 1). Every edge in the pan-network is characterized by its ‘universality number’, U, defined as the total number of our networks in which this edge was observed. More than 80% of the edges were observed in only one network (universality, U=1) suggesting that gene networks in Arabidopsis operating under different conditions are drastically different from each other (Ideker and Krogan, 2012). Co-expression edges between kinases and transcription factors (TF) are often of a particular biological interest as they may indicate gene regulation triggered by signaling pathways. For instance, we observed that a guanylate kinase (AT3G57550) is coexpressed with a MYB TF (AT1G18570) in only one out of the 134 experiments (GSE40354, Supplemental File 1). The experimental context of the coexpression between those two genes involved treatment with bacterial elicitor (Tintor et al., 2013). This suggests further investigation of what exactly makes these proteins so specifically co-regulated. Another edge deserving further investigation connects an F-box protein (AT1G47340) and SKP-1 (AT5G42190) (Supplemental File 1). F-box protein is well known to interact with SKP-1 to degrade unwanted proteins (Schulman et al., 2000), however, hundreds of F-box genes are encoded in the Arabidopsis genome (Kuroda et al., 2002). It is critical to determine the specificity of the interactions between those F-box genes and their interacting partners. It is also important to understand under which environmental conditions the interaction happens (Skaar et al., 2013). The results from our analysis suggest the design of further experiments to reveal the specificity of these interactions. In contrast to the edges that were observed in only one dataset (i.e. U = 1), the edges with larger values of U (Universality) were mostly formed between members of large multi-protein complexes (Supplemental File 1).

### The core-network connects components of large molecular machines

A family of core networks of progressively increasing universality can be extracted from the pan-network by applying a strict cutoff on the universality of edges (e.g. a core-network formed by all edges existing in at least 5 datasets). As the cutoff value increases, the resulting network becomes smaller but more modular (Figure 3). Modularity measures how well are these network modules separated from each other, while the clustering coefficient measures how tightly the neighbors of a node within a module are connected with each other. Both parameters are frequently reported for all types of biological networks, including protein-protein networks, metabolic networks and transcriptional networks (Albert, 2005) but (to the best of our knowledge) ours is the first study of their systematic dependence on edge universality.

**Figure 3.**
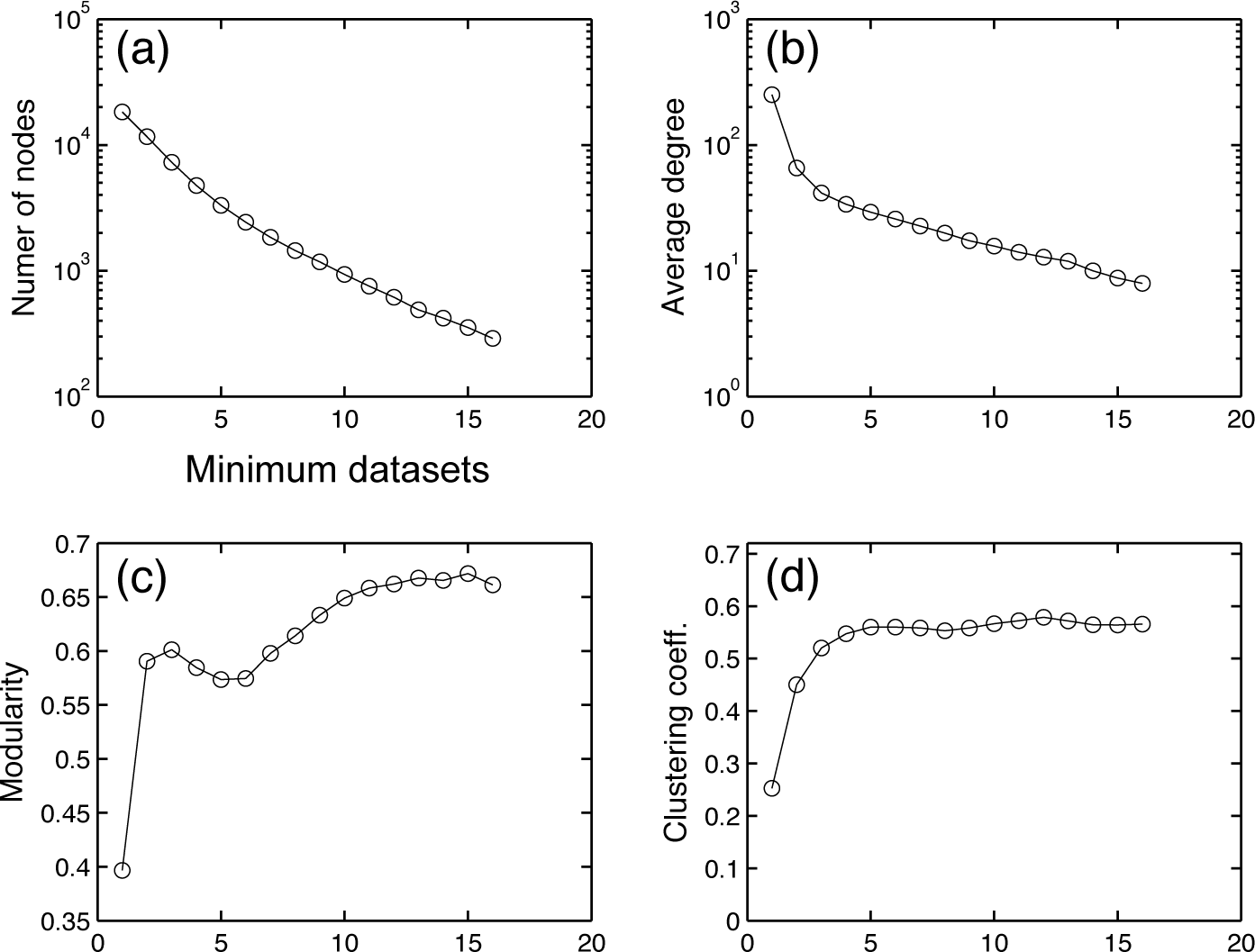
Cutoff selection for construction of the core-network. Edge universality, U_i_, is given by the number of network datasets it was observed. The number of nodes (a), the average degree (b), the modularity (c) and the clustering coefficient (d) of the network constructed from edges with universalities U_i_ greater than or equal to the X-coordinate of the plot.

Based on Figure 3c, we used 10 as the cutoff to determine the core-network used in the rest of our study (marked red in Figure 4), which contains 7326 non-redundant edges among 935 genes. We refer to the set of edges present in the pan-network but not universal enough to be included in the core network as “condition-specific” (marked green in Figure 4). The degree distribution of the core-network approximately follows a power-law (scale-free) pattern with the exponent −1.5 transitioning into exponential cutoff above 50 (Figure 5, Supplemental File 2) (Barabási and Albert, 1999; Barabási and Bonabeau, 2003). Many of its edges connect parts of large multi-protein complexes such as the ribosome and photosystems (Supplemental File 3). In fact, the network is enriched for physically interacting (binding) proteins (Chatr-Aryamontri et al., 2015) (*p-value* < 0.001, Figure 4). The modules of this network were also highly enriched in genes characterized by functional categories ‘translation’ or ‘photosynthesis’ (*p-value* < 10^−10^, Supplemental File 4). These facts are consistent with earlier observation that the genes involved in large molecular machines tend be coexpressed under many conditions (Mao et al., 2009).

**Figure 4.**
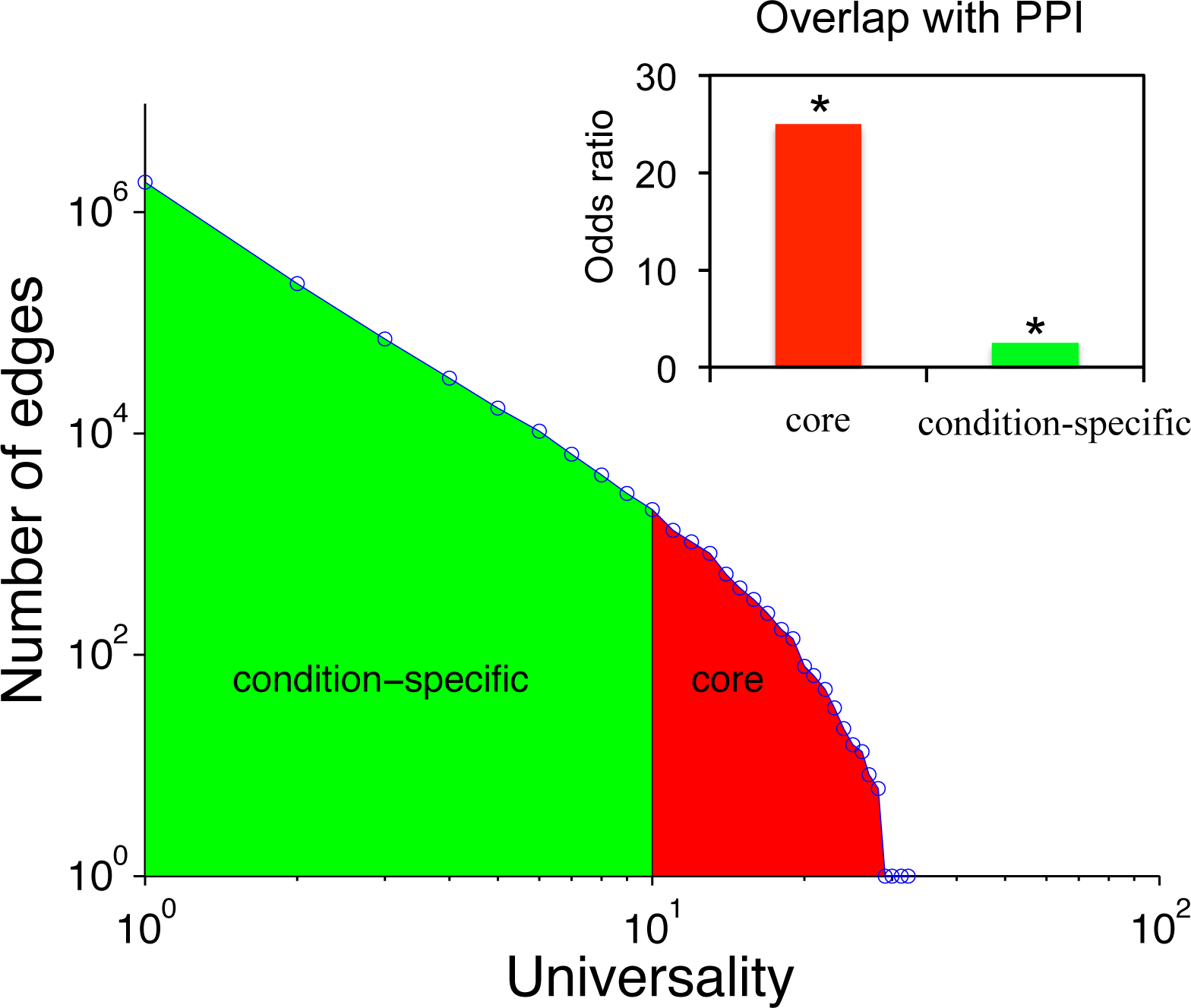
The pan-network contains both core and condition-specific edges. The overlap with the protein-protein interaction (PPI) network for both types of edges are statistically significant (*p-value* < 0.001). The PPI data was downloaded from BioGRID version 3.4.132 (http://thebiogrid.org/).

**Figure 5.**
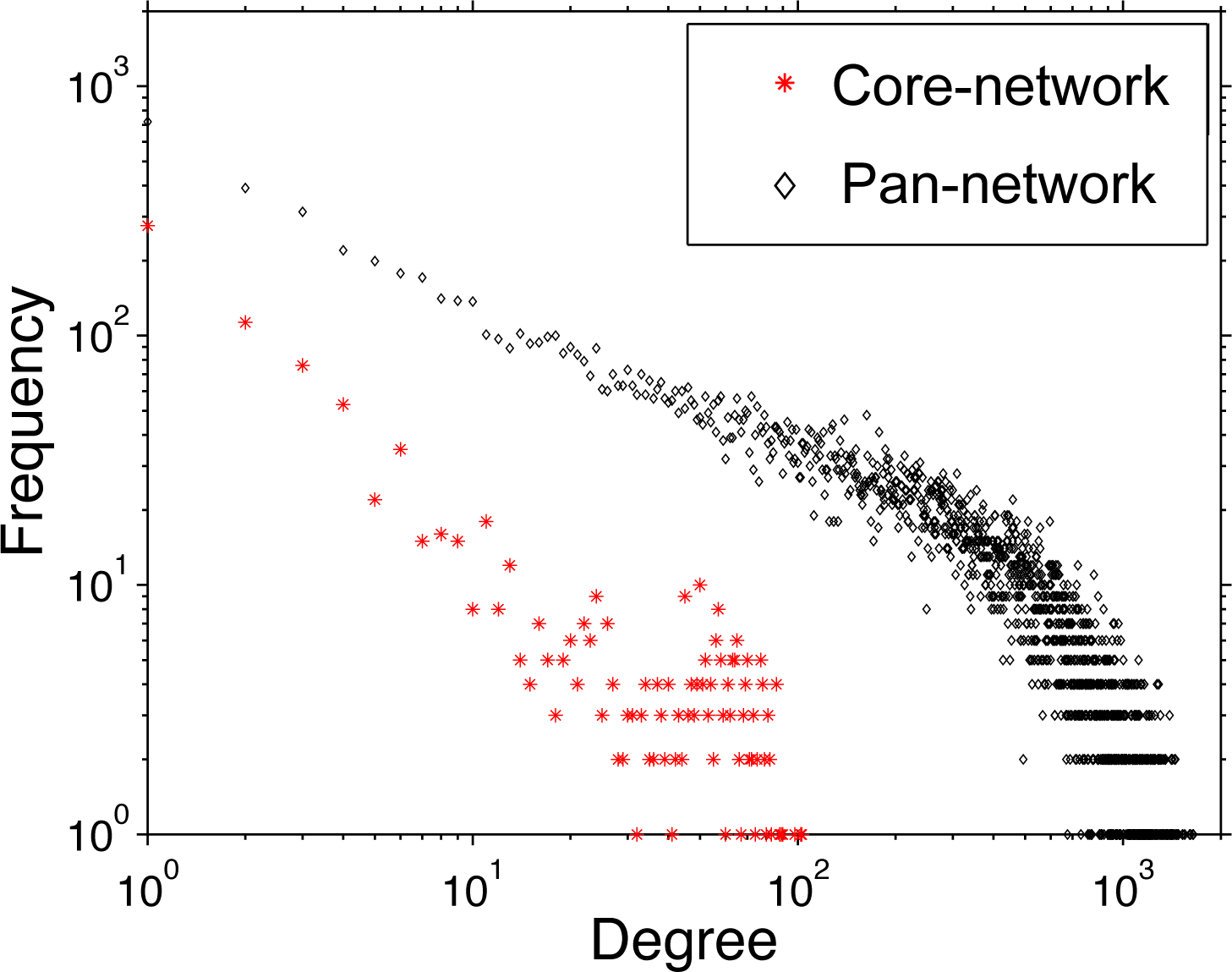
Degree distribution for pan-and core-networks. The y-axis is the number of nodes with a particular degree.

### The pan-network is enriched in condition-specific biological processes

Since each expression dataset usually has its own experimental design addressing a specific biological question (Barrett et al., 2013), an edge detected in one dataset may not be detected in another (de la Fuente, 2010). As more and more evidence supports condition-specific networks in animals (Hudson et al., 2009; Southworth et al., 2009; Anglani et al., 2014), we hypothesized that biological processes that are active in a small set of specific contexts would form the bulk of our pan-network. First of all, although the edges with smaller values of U contained fewer direct physical Protein-Protein Interactions (PPIs) compared with the core-network, PPIs are still overrepresented among pan-network edges (*p-value* < 0.001, Figure 4). This suggests that biologically meaningful connections exist among co-expressed genes which are less likely to be detected by the traditional methods (Lee et al., 2004). The degree distribution for the pan-network approximately followed a power-law pattern (exponent = −0.5 transitioning to the exponential cutoff above 500) (Figure 5, Supplemental File 2). Besides the support from physical interactions, we wondered if more evidence could be found to reveal the biological significance of the pan-network.

With an average degree more than 200 (Supplemental File 2), an overview of the pan-network showed a densely connected large central component. However, we were able to detect network communities (i.e modules) using a scalable algorithm (Lefebvre, 2008) implemented in Gephi 0.9 graph visualization software package (Bastian et al., 2009). In fact, the modules from the core-network and pan-network were identified using the same method to allow for an apples-to-apples comparison. We found two (out of six) modules in the pan-network enriched in ‘regulation of cell cycle’ (*p-value* = 5.39×10^−15^) and ‘regulation of cell communication’ (2.93×10^−6^), respectively (Figure 6). Interestingly, those biological processes were not detected among core-network modules (*p-value* > 0.01, Supplemental File 4).

**Figure 6.**
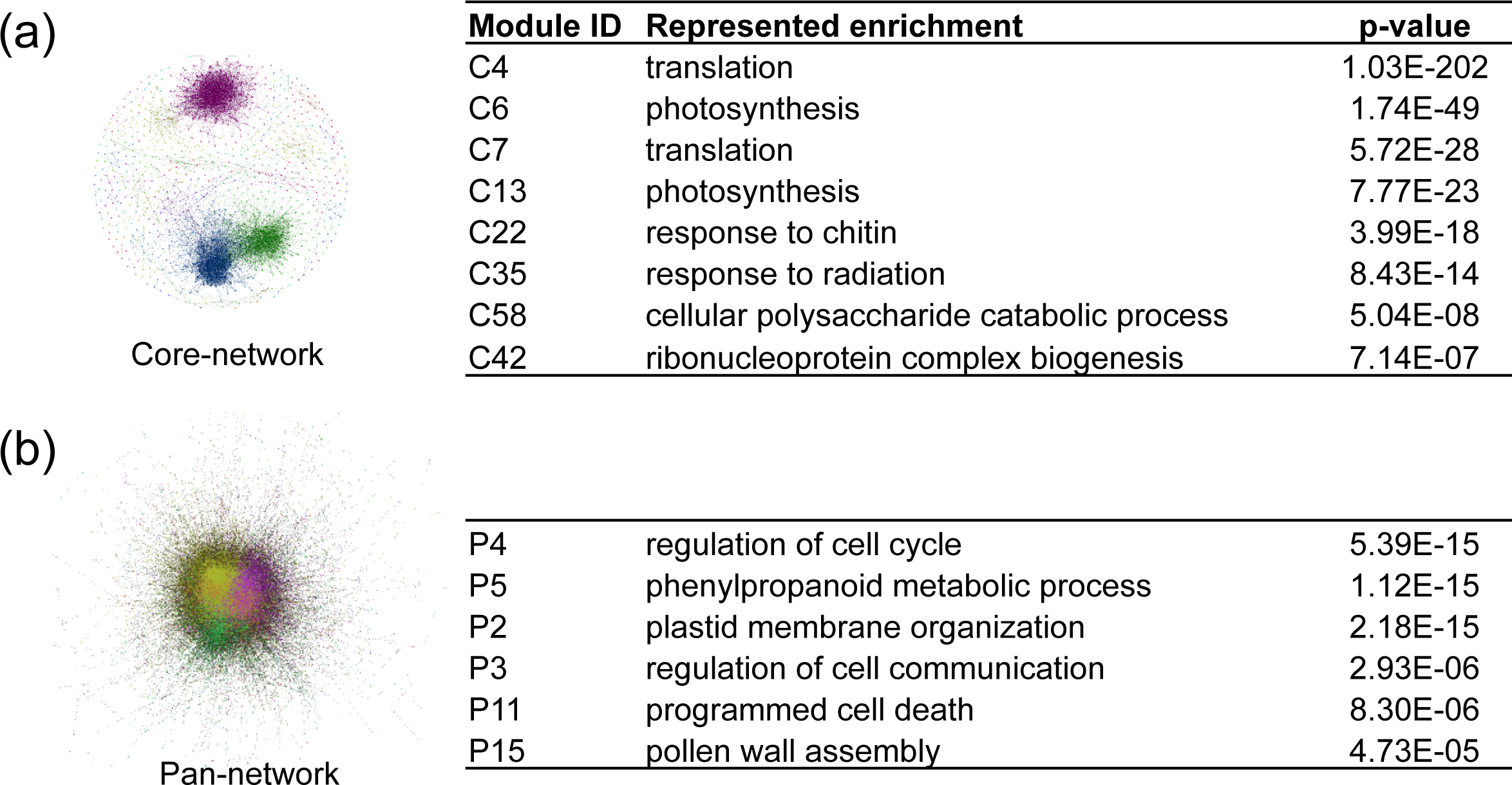
**The functional enrichment for network modules** in pan-network. (a), and in core-network (b). The most over-represented biological processes were shown for each module in the core-network while the biological processes that only enriched in pan-network modules were listed. See Supplemental File 4 for detail.

The most connected nodes, i.e. hubs, are generally the focus of the network analysis. For instance, hubs in protein-protein interaction networks are more likely to be essential genes (Jeong et al., 2001; Zotenko et al., 2008). Hubs in coexpression networks are often considered to be the most informative genes (Horvath and Dong, 2008; Mao et al., 2009; Azuaje, 2014). Approximate power-law degree distribution observed in our analysis confirms the existence of hubs in both pan-network and core-network (Figure 5). Most of the hubs in core-network are ribosomal genes, while the hubs in the pan-network represent a broad spectrum of functional categories, such as aminotransferase (AT3G49680), or ferredoxin (AT1G10960) (Table 1, Supplemental File 2). Genes involved in ‘response to abiotic stimulus’ were enriched among the top 200 most connected genes in the pan-network (*p-value* = 1.6×10^−10^). In addition, chaperonin genes were also enriched (*p-value* < 0.01). Chaperonins play a critical role in helping plants fight against environmental stresses by reestablishing the normal conformation of proteins. This may explain their potential ability to interact with many different genes under different conditions (Wang et al., 2004). For instance, AT1G55490, encoding a subunit of chloroplasts chaperonins, was coexpressed with 1605 genes in the pan-network. These instances of coexpression were from 49 different experiments in total (Supplemental file 5). In conclusion, we demonstrated that a broad spectrum of condition-specific biological processes can be revealed by the pan-network analysis.

**Table1.** The hubs in pan-network and core-network^a^

| **Top 10 hubs in** |  |  |
| --- | --- | --- |
| **pan-network** | **Degree** | **Gene description** |
| AT1G33040 | 1524 | nascent polypeptide-associated complex subunit alpha-like |
| AT2G37660 | 1527 | NAD(P)-binding Rossmann-fold superfamily protein |
| AT1G15820 | 1529 | light harvesting complex photosystem II subunit 6 |
| AT3G49680 | 1549 | branched-chain aminotransferase 3 |
| AT5G46110 | 1551 | Glucose-6-phosphate/phosphate translocator-related |
| AT1G55490 | 1605 | chaperonin 60 beta |
| AT1G74470 | 1607 | Pyridine nucleotide-disulphide oxidoreductase family protein |
| AT3G56940 | 1615 | dicarboxylate diiron protein, putative (Crd1) |
| AT4G01150 | 1642 | located in thylakoid |
| AT1G10960 | 1648 | ferredoxin 1 |
| **Top 10 hubs in** |  |  |
| **core-network** | **Degree** | **Gene description** |
| AT2G46820 | 86 | photosystem I P subunit |
| AT3G60770 | 86 | Ribosomal protein S13/S15 |
| AT4G03280 | 86 | photosynthetic electron transfer C |
| AT4G21280 | 86 | photosystem II subunit QA |
| AT2G36170 | 88 | Ubiquitin supergroup;Ribosomal protein L40e |
| AT1G42970 | 89 | glyceraldehyde-3-phosphate dehydrogenase B subunit |
| AT5G16130 | 90 | Ribosomal protein S7e family protein |
| AT1G26880 | 98 | Ribosomal protein L34e superfamily protein |
| AT1G54780 | 102 | thylakoid lumen 18.3 kDa protein |
| AT3G23390 | 103 | Zinc-binding ribosomal protein family protein |

<sup>a</sup> See Supplemental File 2 for a full list of nodes.

## DISCUSSION

Networks of interactions between different genes are key to our understanding of cellular mechanisms. Large-scale network data for plants are still very limited and are expensive to generate experimentally. Coexpression networks inferred from gene expression profiling data allows one to study interactions between genes (or proteins they encode) albeit indirectly. Based on ‘guilt-by-association’ coexpression network analysis provides great power in predicting gene functions in plants. It also suggests candidates for the design of both high-throughput (Chae et al., 2012; Jiménez-Gómez, 2014) as well as more focused low-throughput experiments (Yonekura-Sakakibara et al., 2007; Mentzen et al., 2008; Pick et al., 2013; Vanholme et al., 2013). Thousands of expression profiling experiments are available for Arabidopsis in the GEO (Barrett et al., 2013). In our previous study we reported principle component analysis of ~7000 expression samples from more than 300 publications in Arabidopsis (He et al., 2016). The developmental stage, growth conditions and the tissues of the mRNA samples used in each study are highly variable, since each of these experiments was designed to answer a specific biological question (Barrett et al., 2013). In this study, we assume the coexpression network inferred from a given experiment represents a part of the overall gene regulatory network perturbed by these specific changes in environmental or intrinsic conditions. We found relatively small number of edges (core network) that can be repeatedly detected in multiple conditions. Differences between functional enrichment within modules in pan-and core-networks emphasize the importance of biological context in coexpression analysis.

Recent studies have shown that coexpression networks in animals are highly condition-specific (Hudson et al., 2009; Southworth et al., 2009; Anglani et al., 2014). Network rewiring was first revealed through comparisons between different cellular states, such as healthy and cancerous tissues (de la Fuente, 2010). Our study showed plant coexpression networks are also under dramatic changes under different conditions. Besides exploring these changes by U of edges, we further calculated the preservation of gene modules in each dataset (Langfelder et al., 2011). Consistent with the results shown in the above, the most preserved modules are enriched in ‘photosynthesis’ (*p-value* < 10^−10^, Supplemental File 6). The modules which can only be detected in one dataset are enriched in more specific biological processes, such as ‘pollen exine formation’ (*p-value* < 10^−10^) and ‘nucleosome organization’ (*p-value* = 3.9*10^−7^) (Supplemental File 6). In the plant community, the context-specificity of gene coexpression has been usually ignored (Gigolashvili et al., 2009; Mao et al., 2009; Gu et al., 2010; Mutwil et al., 2011; Pick et al., 2013). Our search of existing literature revealed that most previous meta-analysis studies of plant expression data (except for a few notable examples discussed below) combined datasets from different labs to detect pairs of universally co-expressed genes (Supplemental File 7). Using 15 rice gene expression datasets, *Childs et al*. compared coexpression networks generated from the combined expression data against those from individual datasets. They found networks from individual datasets to contain specific but potentially informative gene modules (Childs et al., 2011). *Lee et al*. detected coexpression relationships based on individual datasets instead of one combined dataset for Arabidopsis as well as for rice (Lee et al., 2010; Lee et al., 2011). Although *Lee et al*. successfully predicted gene functions based on the individual networks they constructed, the differences between networks from different labs were not systemically evaluated in their study.

Our collection of 134 co-expression networks in Arabidopsis based on individual experimental series in GEO database can be used to answer the question of whether one should combine multiple expression datasets before constructing the co-expression network and if yes, which datasets can be best grouped together. In principle, by combining together similar series one can get the best of both worlds: increased statistical power to detect significantly correlated genes can be gained without losing condition-specific edges. To shed light on this problem we constructed a 134×134 matrix of similarities between our set of networks. The similarity was estimated using two different measures. The first similarity matrix shown in Figure 7a and made available for download as Supplemental File 8 was constructed in the spirit of pan-and core-network analysis. It quantifies the fraction of edges shared between a pair of networks (See Methods). While clusters of similar networks, corresponding mostly to identical tissue types (empirical *p-value* < 10^−5^ based on permutation tests), are visible already in this measure, the contrast between similar and different networks can be made even sharper (Figure 7) if one uses an alternative similarity measure based on shared modules of co-expressed gene across a pair of networks detected by the WGCNA algorithm’s (Langfelder et al., 2011) method ‘modulePreservation’ (Supplemental File 9). We used this similarity measure to construct the ‘network of networks’ connecting pairs of networks with average module similarity score above 10 (Figure 7). Densely interconnected modules in this ‘network of networks’ represent good candidates for series that can be integrated without significant loss of condition-specific edges. We plan to investigate pros and cons of this approach in a follow up study.

**Figure 7.**
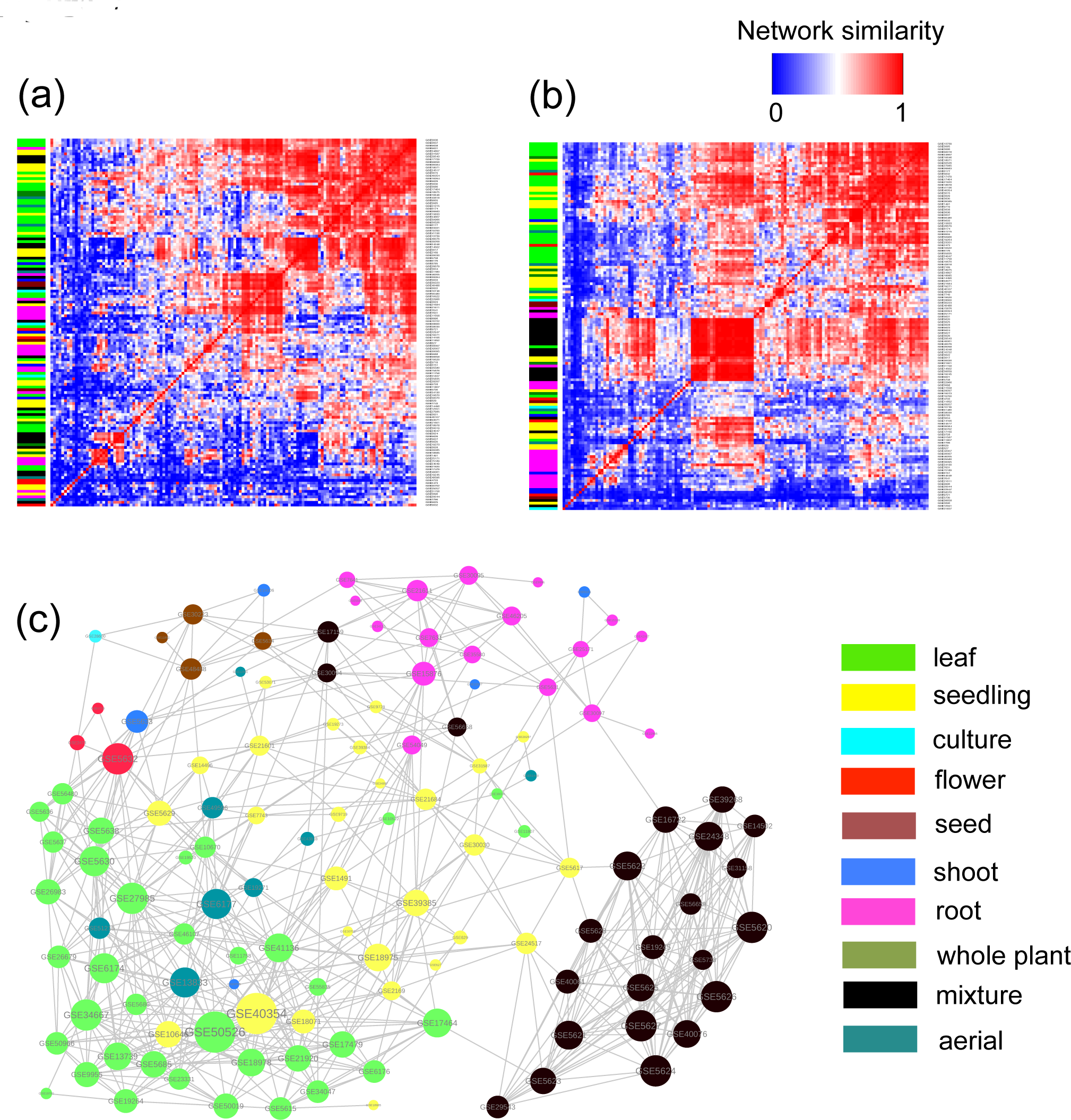
Microarray datasets that are generated from the same tissue type tend to form similar coexpression networks. The clustering of 134 series-based networks based on the similarity between their edges (a) and overlap between modules (b). The ‘network of networks’ (c) connecting pairs of networks with the score of similarity of modules above 10. Network nodes (c) and matrix rows (a, b) are colored according to the tissue type with the color guide shown in the figure.

## CONCLUSION

Our analysis demonstrated that coexpression networks inferred from different microarray datasets share relatively small number of common edges, while at the same time maintaining a large number of condition-specific edges. We constructed a pan-network to represent the union of all detected co-expressed edges among 134 datasets. We also proposed the concept of core-network representing edges detected in multiple datasets. Compared to the pan-network, the core-network is more modular and enriched in genes from large multi-protein complexes. The hubs of the pan-network include genes that play a role in response to a variety of environmental stimuli. In comparison, the modules within the pan-network are enriched in signaling and regulatory functions. We also considered several measures of similarities between individual coexpression networks and constructed ‘network of networks’ connecting similar networks to each other. We anticipate concepts of pan-and core-coexpression networks to provide a useful description of gene regulation architecture in a variety of species and we are currently working on extending our analysis to model organisms other than Arabidopsis.

## METHODS AND MATERIALS

### Microarray data

All the expression datasets in this study were based on the Affymetrix platform GPL198. Only datasets containing at least 20 samples were used (Li et al., 2011). The CEL files of 134 expression datasets were downloaded from GEO (see Supplemental File 10) and normalized by MAS5.0 (Bolstad et al., 2003). The probesets were converted into TAIR gene locus ID based on the annotation file for GPL198. We only used 21678 probesets each of which has a unique mapping on a single Arabidopsis gene.

### Filtering biologically relevant genes

We first applied ANOVA to identify genes that are differentially expressed between replicate groups (*p-value* < 0.01). If there were fewer than 3000 differentially expressed genes, the top 3000 genes ranked by *p-value* were used. We then applied the following standards to exclude genes with low expression. 1) For a gene to be included, at least one of its expression abundance values in a dataset was identified as expressed (i.e. present) by MAS5.0; and 2) For a gene to be included, at least one of its expression abundance values in a dataset was higher than the 90% percentile of the abundance of transposable element on the same array (Rodgers-Melnick et al., 2012). Those biologically relevant genes were then used to build series-based coexpression networks.

### Building the series-based network for each GEO dataset

For each GEO dataset, we calculated the Pearson Correlation Coefficient (PCC) between the expression profiles of two genes. All possible pairs were calculated. Then, the top 0.1% pairs with the highest correlations were used to build the network for each dataset (i.e. *p-value* < 0.001) (Bergmann et al., 2004).

### Enrichment test

The GO annotation data were downloaded from GeneOntology web site(http://geneontology.org/gene-associations/gene_association.tair.gz July 18, 2014). The annotations inferred from the expression profile (i.e. IEP) were removed to avoid the possibility of ‘self-fulfilling prophecy’. All the daughter nodes were recursively mapped to the mother node based on ‘is_a’ relationship. Only the GO terms that are not broadly associated with too many genes were used according to the Bonferroni correction:

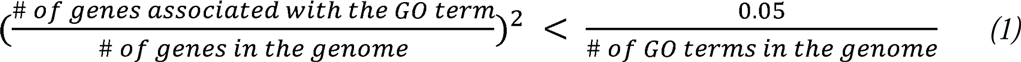

The Fisher’s exact test was utilized to calculate the significance of the enrichment for each GO term, followed by Benjamini-Hochberg correction.

### Calculation of network similarity between different datasets

We first used the fraction of overlapped edges between two networks to measure their similarity (Supplemental File 8) which was based on Jaccard index,

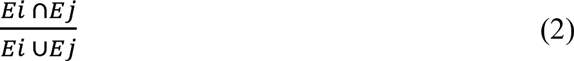

where *Ei* and *Ej* are the edges in network i and j, respectively. Another measure of network similarity was calculated as follows (Supplemental File 9). First, densely interconnected network modules were detected by Weighted Gene Co-Expression Network Analysis (WGCNA) software for each one of our 134 networks (Langfelder and Horvath, 2008). Second, the method, ‘modulePreservation’ within WGCNA was utilized to calculate the preservation of each module in another dataset (Langfelder et al., 2011). Then the average Z-summary score of all the modules shared between a pair of networks was used to represent their similarity. This score was normalized between 0 and 1 for visualization purpose by,

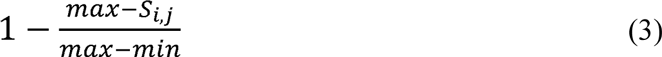

where *S*_*i,j*_ is the similarity (i.e average Z-summary) between a pair of datasets. *max* and *min* represent the largest and smallest similarity, respectively. A cutoff of *S*_*i,j*_ > 10 was applied in order to keep the network pairs with the strongest similarity when be visualized (Langfelder et al., 2011). If a network has more than 10 neighbors, only the first 10 neighbors were shown. If a network has no neighbor, it was not shown. For more information on the calculation of network similarity using WGCNA, see Supplemental File 11.

## SUPPLEMENTAL MATERIALS

### Supplemental File 1

The detailed information for the subset of edges in the pan-network overlapping with the Protein-Protein Interaction network. The entire list of pan-network edges can be downloaded at the BitBucket (URL included in the file).

### Supplemental File 2

Degrees and module memberships of all nodes in core-and pan-networks.

### Supplemental File 3

The table of all edges of the core-network and their degree of universality (the number of networks they were observed).

### Supplemental File 4

The enriched biological processes among network modules for pan-and core-networks.

### Supplemental File 5

The detailed information for the gene AT1G55490 in the pan-network.

### Supplemental File 6

The functional enrichment and preservation of gene modules identified in individual GEO dataset.

### Supplemental File 7

The comprehensive list of publications reporting plant coexpression network meta-analysis along with the information on whether or not the study combines multiple datasets.

### Supplemental File 8

Network similarity table measured by the fraction of shared edges (Eq. 2).

### Supplemental File 9

Network similarity measured by the overlap between shared modules (Eq. 3).

### Supplemental File 10

The table of GEO datasets used in this study.

### Supplemental File 11

Our choice of parameters for the WGCNA software package including its ‘modulePreservation’ method.

## ACKNOWLEDGEMENTS

We thank Shinjae Yoo, Daifeng Wang, Mark Gerstein, Sunita Kumari and Doreen Ware for helpful discussions. We also appreciate editing provided by Claudia Lutz from the University of Illinois Urbana-Champaign.

